# PI3 kinase inhibitor PI828 uncouples aminergic GPCRs and Ca^2+^ mobilization irrespectively of its primary target

**DOI:** 10.1101/2024.01.17.576005

**Authors:** Polina D. Kotova, Ekaterina A. Dymova, Oleg O. Lyamin, Olga A. Rogachevskaja, Stanislav S. Kolesnikov

## Abstract

The phosphoinositide 3-kinase (PI3K) is involved in regulation of multiple intracellular processes. Although the inhibitory analysis is generally employed for validating a physiological role of PI3K, increasing body of evidence suggests that PI3K inhibitors can exhibit PI3K-unrelated activity as well. Here we studied Ca^2+^ signaling initiated by aminergic agonists in a variety of different cells and analyzed effects of the PI3K inhibitor PI828 on cell responsiveness. It turned out that PI828 inhibited Ca^2+^ transients elicited by acetylcholine (ACh), histamine, and serotonin, but did not affect Ca^2+^ responses to norepinephrine and ATP. Another PI3K inhibitor wortmannin negligibly affected Ca^2+^ signaling initiated by any one of the tested agonists. Using the genetically encoded PIP_3_ sensor PH(Akt)-Venus, we confirmed that both PI828 and wortmannin effectively inhibited PI3K and ascertained that this kinase negligibly contributed to ACh transduction. These findings suggested that PI828 inhibited Ca^2+^ responses to aminergic agonists tested, involving an unknown cellular mechanism unrelated to the PI3K inhibition. Complementary physiological experiments provided evidence that PI828 could inhibit Ca^2+^ signals induced by certain agonists, by acting extracellularly, presumably, through their surface receptors. For the muscarinic M3 receptor, this possibility was verified with molecular docking and molecular dynamics. As demonstrated with these tools, wortmannin could be bound in the extracellular vestibule at the muscarinic M3 receptor but this did not preclude binding of ACh to the M3 receptor followed by its activation. In contrast, PI828 could sterically block the passage of ACh into the allosteric site, preventing activation of the muscarinic M3 receptor.

## 1. Introduction

The superfamily of G-protein coupled receptors (GPCRs) includes the subgroup of aminergic receptors for biogenic amines such as dopamine, epinephrine, norepinephrine, histamine, and serotonin. Based on molecular structure, muscarinic acetylcholine (ACh) receptors have also been included in this GPCR subfamily (Vass et al., 2019). The aminergic GPCRs are ubiquitously expressed throughout the body and involved in the regulation of various physiological processes from neurotransmission and muscle contraction to cell growth and differentiation (Archer et al., 2021; Ledonne and Mercuri, 2017; Obara et al., 2020; van der Westhuizen et al., 2020; Wu et al., 2019). The capability of aminergic GPCRs to control so diverse physiological functions is provided by their coupling to a variety of downstream effectors and signaling pathways, including adenylyl cyclase, phospholipase C (PLC), small G-proteins, NO synthase, ion channels, and various kinases, particularly, class I phosphoinositide 3-kinases (PI3Ks) (Nakano et al., 2017).

PI3Ks phosphorylate phosphatidylinositol (3,4)-bisphosphate (PIP_2_) in the plasma membrane, producing phosphatidylinositol (3,4,5)-trisphosphate (PIP_3_) and stimulating PIP_3_-dependent intracellular processes (Cantley, 2002; Vanhaesebroeck et al., 2012). PIP_3_ carries out its signaling function primarily through the recruitment of intracellular proteins with PIP_3_-binding domains and motifs, including the pleckstrin homology (PH) domain (DiNitto and Lambright, 2006). Notable among PH domain-containing proteins are the kinases PDK1 and Akt (Jean and Kiger, 2014; Vanhaesebroeck et al., 2010), which are involved in regulation of multiple intracellular targets in a PIP_3_-dependent manner. In particular, the PI3K/Akt pathway can modulate Ca^2+^ release by controlling activity of IP_3_ receptors and PLC (Frégeau et al., 2011; Szado et al., 2008; Zhang et al., 2009). The involvement of the PI3K/Akt axis in the regulation of Ca^2+^ release was particularly demonstrated in MDCK cells (Santoso et al., 2011), COS-7 cells (Marchi et al., 2012), and RINm5F cells (Frégeau et al., 2011). The PI3K/Akt pathway has also been documented as an important regulator of intracellular Ca^2+^ signaling and excitation-contraction coupling in cardiomyocytes (Ghigo et al., 2017; Graves et al., 2012).

The inhibitory analysis is universally employed as a quite effective approach for uncovering physiological roles of intracellular signaling circuits and regulatory molecules, including PI3K. The disadvantage of this method is that many, if not all, available modulators, antagonists, and inhibitors are not entirely specific toward a single target but can affect several unrelated cellular proteins. This also is the case with PI3K inhibitors, including wortmannin and LY294002, which are widely used to validate a contribution of the PI3K-based pathways to intracellular signaling. It has particularly been demonstrated that LY294002 not only bound to different classes of PI3K, but also bound to a wide range of other proteins (Gharbi et al., 2007) and was capable of inhibiting a number of protein kinases, such as mammalian target of rapamycin (mTOR) (Brunn et al., 1996) and DNA-dependent protein kinase (Smith et al., 1999), casein kinase-2 (CK2) (Davies et al., 2000), Pim-1 kinase (Jacobs et al., 2005). The list of effects, unrelated to the PI3K inhibition, exerted by LY294002 also includes the inhibition of nuclear factor-κB (Choi et al., 2004; Kim et al., 2005) and induction of expression of transcription factor 3 (Yamaguchi et al., 2006), the inhibition of multidrug resistance-associated protein MRP1 (Abdul-Ghani et al., 2006) and the BET bromodomain proteins (Dittmann et al., 2014) as well as the significant enhancement of intracellular H_2_O_2_ production (Poh and Pervaiz, 2005). Nonspecific mechanisms can mediate inhibitory effects of LY294002 on intracellular Ca^2+^ signaling as well (Ethier and Madison, 2002; Kotova et al., 2020; Tolloczko et al., 2004).

The compound PI828 is a LY294002 analogue and more potent PI3K antagonist (Gharbi et al., 2007). Here we studied agonist-induced Ca^2+^ signaling in a variety of different cells and found PI828 to inhibit cell responsiveness to aminergic agonists. Based on physiological results and data obtained with methods of computational biophysics, we inferred that PI828 suppressed agonist-induced Ca^2+^ signaling by acting extracellularly as an antagonist of aminergic GPCRs.

## 2. Materials and methods

### 2.1. Cell lines and culturing

In this study, four cell lines were employed, including HEK-293, С6, and two monoclonal cell lines HEK/R-GECO1/PH(Akt)-Venus and CHO/5-HT_2C_. While HEK-293 and HEK/R-GECO1/PH(Akt)-Venus were cultured in the Dulbecco’s modified Eagle’s medium (DMEM; Invitrogen), С6 and CHO/5-HT_2C_ were maintained in the F12 medium (Invitrogen). In all cases, the culture medium contained 10% (vol/vol) fetal bovine serum (HyClone), glutamine (1%) and the antibiotic gentamicin (100 μg/ml) (Invitrogen). Cells were grown in 12-well culture plates at 37 °C in a humidified atmosphere of 5% CO_2_.

### 2.2. Monoclonal cell lines

The CHO/5-HT_2C_ line, which was derived from CHO-K1 cells, was a monoclonal cell line that stably expressed serotonin 5-HT_2C_ receptors cloned from the mouse brain (Cherkashin et al., 2022).

The monoclonal cell line HEK/R-GECO1/PH(Akt)-Venus, which stably expressed sensor for cytosolic Ca^2+^ R-GECO1 and sensor for PIP_3_ PH(Akt)-Venus, was generated as follows. HEK-293 cells were transfected with two plasmid vectors CMV–R-GECO1 (a gift from Robert Campbell, Addgene plasmid #32444; http://n2t.net/addgene:32444; RRID:Addgene_32444) and PH(Akt)-Venus (a gift from Narasimhan Gautam, Addgene plasmid #85223; http://n2t.net/addgene:85223; RRID:Addgene_85223) (O’Neill and Gautam, 2014; Zhao et al., 2011). Before the day of transfection, cells were put in a 12-well culture plate at the density of 2 – 5 × 10^5^ cells. The transfection mixture (0.5 μg CMV–R-GECO1, 0.5 μg PH(Akt)-Venus, and 4 μl of FuGENE 6 (Promega) in 100 μl of OPTI-DMEM) was added to the cells for transfection. After 24 h incubation, the transfection mixture was replaced with the growth culture medium, and 48 h after transfection, cells were transferred into a 60 mm petri dish. Next, cells were maintained in the growth medium supplemented with 0.6 mg/ml G-418 for 3 weeks. The antibiotic-resistant cells were distinguished by R-GECO1 (ex = 561 nm, em = 610 ± 10 nm) and PH(Akt)-Venus (ex = 488 nm, em = 515 ± 10 nm) emission using a FACSAria SORP cell sorter (BD Biosciences), and those exhibiting most intensive fluorescence were collected individually for further culturing in a 96-well plate. The single cell-derived colonies were grown in the presence of 0.3 mg/ml G-418 for 80% confluence. Specific functionality of each cellular monoclone was examined with Ca^2+^ and PIP_3_ imaging, and a cell line with highest responsivity to ACh and insulin was selected for further experimentation.

### 2.3. Ca^2+^ imaging

For cell isolation, cultured cells were washed with the Versene solution (Sigma-Aldrich), incubated in the 0.25% Trypsin-EDTA solution (Sigma-Aldrich), resuspended in a complete cell growth medium. Isolated cells were plated onto a hand-made photometric chamber of nearly 150 μL volume. The chamber was a disposable coverslip (Menzel-Glaser) with an attached ellipsoidal resin wall; the chamber bottom was coated with Cell-Tak (Corning) ensuring sufficient cell adhesion. Attached cells were loaded with the Ca^2+^ dye Fluo-8 at room temperature (23 – 25 °C) by adding 4 μM Fluo-8 AM (AAT Bioquest) and 0.02% Pluronic F-127 (SiChem) to the bath. Cells were loaded for 20 min, rinsed several times, and kept in the bath solution for 1 h prior to recordings. The bath solution contained (mM): 130 NaCl, 5 KCl, 2 CaCl_2_, 1 MgCl_2_, 10 glucose, 10 HEPES (pH 7.4). When necessary, 2 mM CaCl_2_ in the bath was replaced with 0.5 mM EGTA + 0.4 mM CaCl_2_, thus reducing free Ca^2+^ to nearly 260 nM at 23 °С as calculated with the Maxchelator program (http://maxchelator.stanford.edu). The used salts and buffers were from Sigma-Aldrich.

Experiments were carried out using an inverted fluorescent microscope Axiovert 135 equipped with an objective Plan NeoFluar 20x/0.75 (Carl Zeiss), a digital EMCCD camera Luca^EM^ R 604 (Andor Technology), and a hand-made computer-controllable epi-illuminator with a set of light-emitting diodes that enabled multiwave excitation. Fluo-8 was excited at 480 ± 10 nm, and its emission was collected at 535 ± 25 nm. Serial fluorescent images were captured every second and analyzed using imaging software NIS-Elements (Nikon). Deviations of cytosolic Ca^2+^ from the resting level in individual Fluo-8-loaded cells were quantified by the ratio ΔF/F_0_, where ΔF = F − F_0_, F is the instant intensity of cell fluorescence, F_0_ is the intensity of cell fluorescence recorded in the very beginning of a recording and averaged over a 20 s interval. For data and graphical analysis Sigma Plot 12.5 (Systat Software Inc) was used. “n” represents the number of cells from at least three experiments conducted on different days. All chemicals were applied by the complete replacement of the bath solution in a 150 μl photometric chamber for nearly 2 s using a perfusion system driven by gravity.

### 2.4. PIP_3_ and Ca^2+^ imaging

HEK/R-GECO1/PH(Akt)-Venus cells were cultured in a hand-made photometric chamber for 24 h prior to experiments. The chamber was a Plexiglas framework with an ellipsoidal slot of nearly 150 μl volume, the bottom of the chamber was a disposable coverslip (Menzel-Glaser). Experiments were carried out using an inverted fluorescent microscope Axiovert 200 equipped with an objective Plan NeoFluar 20x/0.75 (Carl Zeiss), a digital sCMOS camera Zyla 4.2P (Andor Technology), metal halide light source AMH-200-F6S (Andor Technology), and spinning disk for confocal microscopy Revolution DSD2 (Andor Technology) (Manton, 2022). Fluorescence of R-GECO1 was excited at 560 ± 20 nm, emission was collected at 630 ± 37 nm. Fluorescence of PH(Akt)-Venus was excited at 500 ± 10 nm, emission was collected at 535 ± 15 nm. Serial fluorescent images were captured with NIS-Elements, depending on an experimental protocol. All chemicals were applied by the complete replacement of the bath solution in a 150 μl photometric chamber for nearly 2 s using a perfusion system driven by gravity.

### 2.5. Drugs

The used ACh, histamine, serotonin, insulin, PI828, U73122, and 4-DAMP were purchased from Tocris Bioscience; norepinephrine was from Calbiochem Biochemicals; ATP and wortmannin were from Sigma-Aldrich; hematein and ketanserin were from MedChemExpress.

### 2.6. Structure of the muscarinic M3 receptor

Given that a 3D-structure of the human muscarinic M3 receptor is currently not available, while its amino-acid sequence is more than 90% homologous to the rat M3 receptor, PDB ID: 4U15 (Thorsen et al., 2014), the crystal structure of the last was used as a template. The original structure obtained from RCSB crystallographic database https://www.rcsb.org (Berman et al., 2000) was modified by removing manually all non-protein molecules, e.g. the antagonist tiotropium, from the initial structure, and also a protein segment between the 5th and 6th transmembrane helices, which has been added to facilitate the crystallization. The resultant gap at the intracellular loop 3 was filled in with the amino acid linker GSGSGSGS using the SWISS-MODEL tool (Waterhouse et al., 2018). This sequence of glycine and serine residues was chosen taking into account their common occurrence in natural linkers, electroneutrality, and short side chains. In addition, similar sequences were previously used for modeling ligand-receptor interactions (Sterling and Irwin, 2015). The number of chosen GS-repeats provided the necessary minimum that allowed for unconstrained dynamics of the 5th and the 6th helices of the M3 receptor. Finally, the homologous modeling of human M3 receptor structure was accomplished using the SWISS-MODEL tool, based on the rat M3 receptor template described above.

### 2.7. Molecular docking

3D-structures of 4-DAMP, wortmannin, and PI828 were obtained from the databases ZINC (Sterling and Irwin, 2015) and Pubchem (Kim et al., 2021). All these structures were prepared for docking with AutoDockTools-1.5.6. Docking was performed with Audock Vina 1.1.2 (Trott and Olson, 2010). This tool allowed for constraining the search space of conformations energetically most favorable for forming a stable ligand-receptor complex by specifying an appropriate region on the receptor surface. The region between M3 receptor transmembrane helices, which was virtually bounded by their extracellular loops and the receptor center, appeared to be rather suitable, given a number of experimentally-obtained crystal structures of muscarinic receptors in complexes with their known antagonists (e.g. with tiotropium, PDB ID: 4U15).

Since no 3D-structure of the ACh-M3 receptor complex is available now, the location of ACh on the receptor was modeled similarly with docking and was further considered as the orthosteric ACh-binding site.

### 2.8. Molecular dynamics

To eliminate steric clashes potentially produced by the docking, refine ligand pose on the receptor surface, and validate stability of the docking-predicted complexes, we performed molecular dynamics simulations using the pmemd.cuda tool, which is included in the Amber Molecular Dynamics Package (version 20) (Case et al., 2021), and GPUs Nvidia 2080 and Nvidia 3080.

Protonation states of the M3 receptor amino-acid residues were preliminarily determined. Protonation states of all residues were predicted by PROPKA at pH 7.0 (Jurrus et al., 2018), except for Asp 113 which was protonated. The conserved Asp 113 corresponds to Asp 83 in rhodopsin, which is known to be protonated during the entire photocycle (Fahmy et al., 1993). Amber preparation tools charge terminal residues by default. To avoid that, N- and C-termini were capped with neutral acetyl and N-methylamide groups, correspondingly.

The structure was then oriented with OPM (Lomize et al., 2012) for packing in a membrane and inserted into a POPC bilayer and solvated using Packmol Memgen 1.1.0 (Martínez et al., 2009; Nugent and Jones, 2013). To neutralize the net charge, sodium and chloride ions were added at the concentration of 150 mM. The final molecular system contained 100000 atoms in the nearly 100 × 100 × 100 Å volume.

The Amber force field parameters were adjusted to the structures as follows: ff19SB – for proteins, lipid17 – for lipids, gaff2 – for ligands. The four-point OPC water model was used. Following 5000-step minimization done with the Amber package tool pmemd, the system was heated to 310 K and equilibrated in two stages: under the canonical NVT ensemble for 1 ns, first, and under the isothermal-isobaric NPT ensemble at 1 atm for 1 ns, afterwards. Production simulations at 310 K and 1 atm under the NPT ensemble were then initiated and run for at least 100 ns. Covalent bonds involving hydrogen atoms were constrained using SHAKE. Short-range non-bonded interactions were cut off at 9 Å. The Langevin thermostat and the Berendsen (equilibration) and the Monte-Carlo (production runs) barostats were used. Long-range electrostatic interactions were computed using the Particle Mesh Ewald method.

Accelerated molecular dynamics modeling was performed under the same conditions according to the algorithm described in the Amber manual. Said algorithm speeds up conformational transitions by reducing the energy barriers between the states of the simulated system. Energy threshold and boosting factor parameters were calculated with the formulas presented in the manual.

Publication-quality images were generated in PyMOL (Schrodinger, 2015).

## 3. Results

### 3.1. Effects of PI3K inhibitors on Ca^2+^ signaling initiated by aminergic agonists

The experiments described below were performed using cells of several lines. ACh and norepinephrine-induced Ca^2+^ signaling was studied in HEK-293 cells known to express endogenous muscarinic and adrenergic receptors (Atwood et al., 2011). Cells of rat glioma (C6) responded to histamine with Ca^2+^ transients as well. Serotonin-induced Ca^2+^ signaling were studied in CHO/5-HT_2C_ cells that stably expressed the recombinant serotonin 5-HT_2C_ receptor (Cherkashin et al., 2022). Given that cells of the above-mentioned lines were not responsive, in terms of Ca^2+^ signaling, to dopamine, dopaminergic signaling was not studied here. In a typical experiment, Fluo-8 loaded cells were shortly (∼ 100 s) stimulated by a particular agonist at moderate concentrations every ∼300 s, the period being sufficient for post-stimulation recovery of cells and prolongation of their responsiveness during long-lasting recordings.

The pulse stimulation of HEK-293, C6, and CHO/5-HT_2C_ cells with ACh (1147 cells), histamine (353 cells), and serotonin (508 cells), respectively, triggered Ca^2+^ transients in control but not in the presence of PI828; the inhibitory effects of which were reversible (Fig. 1A–C). Nearly half cells, which responded to 1 μM ACh, 2 μM histamine, and 1 nM serotonin, became unresponsive to the agonists with PI828 in the bath applied at 20 μM, 20 μM, and 10 μM (∼IC_50_), respectively. In contrast, at the same doses, PI828 canceled neither norepinephrine (3 μM) responses of HEK-293 cells (*n* = 122) (Fig. 1D) nor ATP (3 – 5 μM) responses of cells of all assayed types (Fig. 1A–C).

**Fig. 1.**
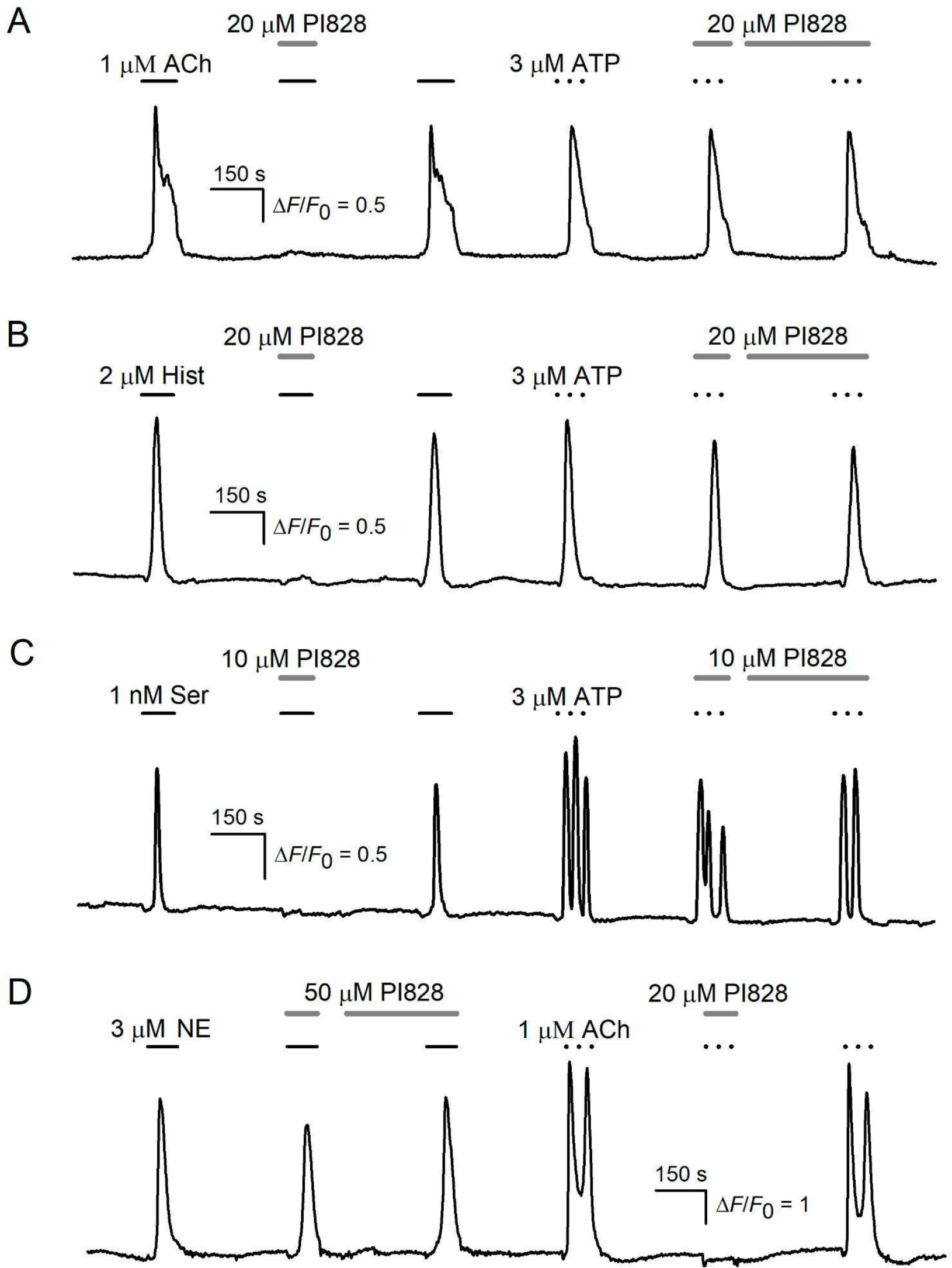
Effects of PI828 on cells responsiveness to aminergic agonists. Representative recordings of intracellular Ca^2+^ in individual Fluo-8-loaded cells stimulated by aminergic agonists in control and in the presence of PI828. PI828 (10 – 50 μM) inhibited Ca^2+^ transients elicited by 1 μM ACh in HEK-293 cells (A), 2 μM histamine in C6 cells (B), and 1 nM serotonin in CHO/5HT_2C_ cells (C). Note that PI828 did not inhibit cell responsiveness to the purinergic agonist ATP (3 μM) (A–C). (D) Representative recording from a HEK-293 cell demonstrating that unlike ACh responses, Ca^2+^ responses to 3 μM norepinephrine (NE) were not inhibited by 50 μM PI828. Here and in the below figures: applications of compounds are indicated by the straight-line segments above the experimental traces; the data are presented as ΔF/F_0_, where ΔF = F–F_0_, F is the instant intensity of cell fluorescence, F_0_ is the intensity of cell fluorescence obtained in the very beginning of a recording and averaged over a 20 s interval.

The inhibitory effects of PI828 on cellular responses to ACh, histamine, and serotonin could formally be considered as evidence for the involvement of PI3K in the transduction of these aminergic agonists. However, when cells were treated, as internal control, with another PI3K inhibitor wortmannin (1 – 10 μM), this compound did not affect the responsiveness of assayed cells to ACh (*n* = 154), histamine (*n* = 127), and serotonin (*n* = 176) (Fig. 2). These findings summarized in Table 1 questioned whether the observed inhibitory effects of PI828 on cell responsiveness (Fig. 1 and Fig. 2) were specifically associated with the suppression of PI3K activity.

**Fig. 2.**
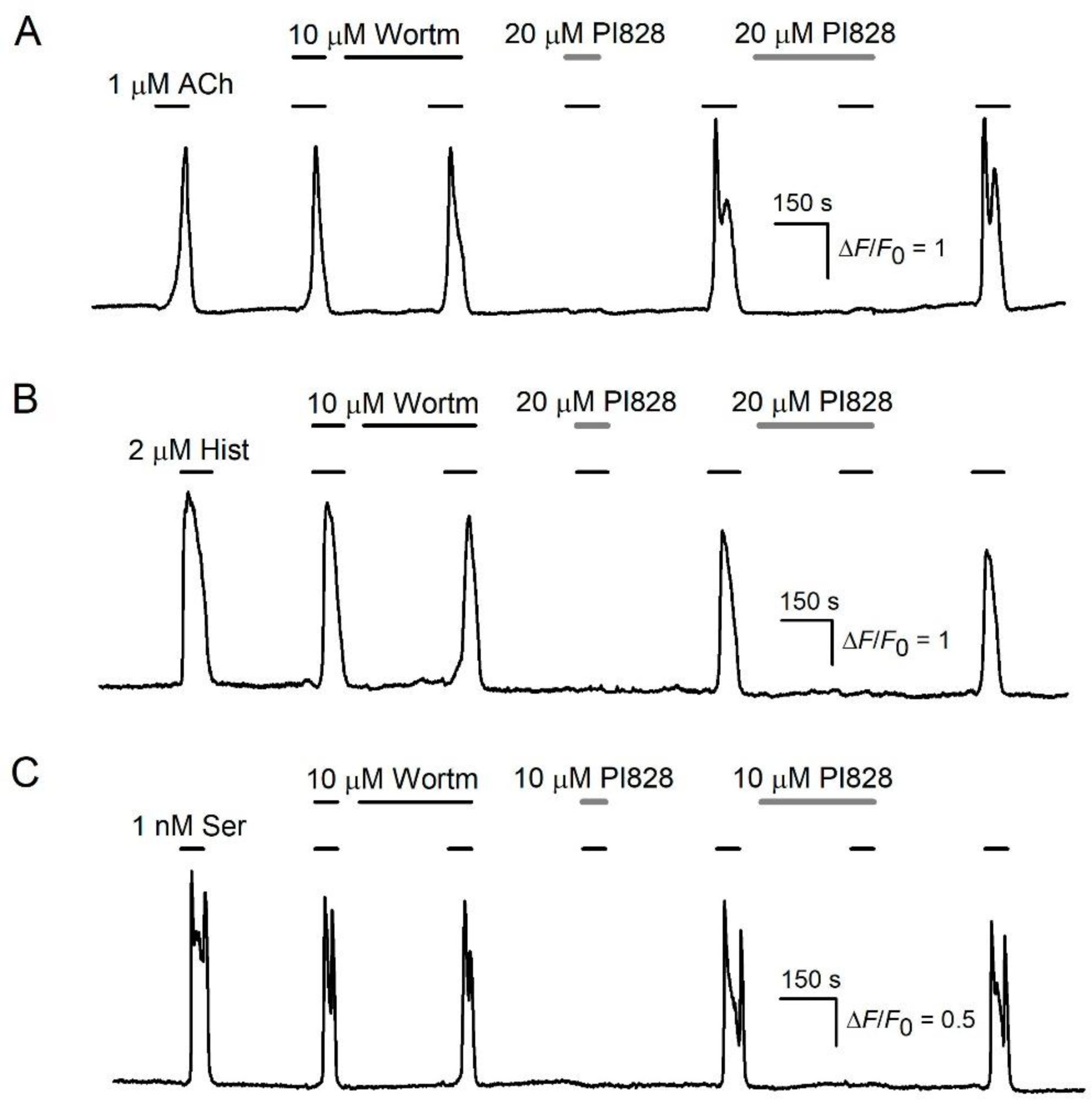
Dissimilar effects of wortmannin and PI828 on cells responsiveness to aminergic agonists. (A–C) Representative responses of cells to aminergic agonists in control and in the presence of wortmannin or PI828. Unlike PI828 (10 – 20 μM), wortmannin (10 μM) did not affect Ca^2+^ transients elicited by 1 μM ACh in HEK-293 cells (A), 2 μM histamine in C6 cells (B), and 1 nM serotonin in CHO/5HT_2C_ cells (C).

**Table 1.** Effects of PI828 and wortmannin on cell responsiveness to agonists.

| Agonist | Effect of PI828 | Effect of wortmannin | Cell line |
| --- | --- | --- | --- |
| Acetylcholine | inhibition, $IC_{50} \sim 20 \mu M$ | no effect at 10 $\mu M$ | HEK-293 |
| Histamine | inhibition, $IC_{50} \sim 20 \mu M$ | no effect at 10 $\mu M$ | C6 |
| Serotonin | inhibition, $IC_{50} \sim 10 \mu M$ | no effect at 10 $\mu M$ | CHO/5HT<sub>2C</sub> |
| Norepinephrine | no effect at up to 50 $\mu M$ | no effect at 10 $\mu M$ | HEK-293 |
| ATP | no effect at up to 50 $\mu M$ | no effect at 10 $\mu M$ | all mentioned |

### 3.2. Inhibition of PI3K activity by wortmannin and PI828

The different effects of the PI3K inhibitors on cell responsiveness to ACh, histamine, and serotonin (Fig. 2) could be attributed to that at concentrations used, PI828 suppressed PI3K isoforms specific for assayed cells much more effectively than wortmannin. We therefore attempted to evaluate specific activity of PI828 and wortmannin as PI3K inhibitors by using HEK/R-GECO1/PH(Akt)-Venus cells, which stably expressed genetically encoded sensors for cytosolic Ca^2+^ (R-GECO1) and PIP_3_ (PH(Akt)-Venus). The R-GECO1 is the fluorescent Ca^2+^ sensor, so that fluorescence of R-GECO1-positive cells increases as cytosolic Ca^2+^ rises (Zhao et al., 2011). The PH(Akt)-Venus sensor operates in another manner. While the total fluorescence of a PH(Akt)-Venus expressing cell is apparently independent of a PIP_3_ level, stimulus-elicited generation of PIP_3_ in the plasmalemma stimulates translocation of PH(Akt)-Venus from the cytosol to plasma membrane (O’Neill and Gautam, 2014). This phenomenon could be clearly visualized with high-resolution microscopy, including structured illumination microscopy (SIM-microscopy) employed in this study.

To initiate PIP_3_ generation, we used insulin known to stimulate tyrosine kinase receptors coupled particularly to the PI3K/Akt pathway (Hopkins et al., 2020). The treatment of HEK/R-GECO1/PH(Akt)-Venus cells with insulin (100 nM) expectedly led to the translocation of PH(Akt)-Venus molecules from the cytosol to the plasmalemma for 3 – 5 min after the insulin application. This effect of insulin was reversible, the 10 min rinse of the assayed cells with the insulin-free bath solution initiated the relocation of PH(Akt)-Venus to the cytosol. Based on these findings and using the protocol shown on Fig. 3A, we examined the activity of wortmannin and PI828 as PI3K inhibitors. It turned out that wortmannin (10 μM) prevented the insulin-induced translocation of PH(Akt)-Venus to the plasmalemma (Fig. 3B) (*n* = 524). When cells were rinsed out for nearly 10 min to remove this PI3K inhibitor, insulin applied once again was capable of initiating the PH(Akt)-Venus translocation, which was indicated by the appearance of relatively bright and contrast narrow zones corresponding to plasma membranes (Fig. 3B, right panel). In the similar assay, PI828 (30 μM) also reversibly inhibited insulin-induced translocation of PH(Akt)-Venus fluorescence (Fig. 3C) (*n* = 438). These observations suggested that both wortmannin and PI828 effectively inhibited PI3K isoforms operating in HEK-293 cells.

**Fig. 3.**
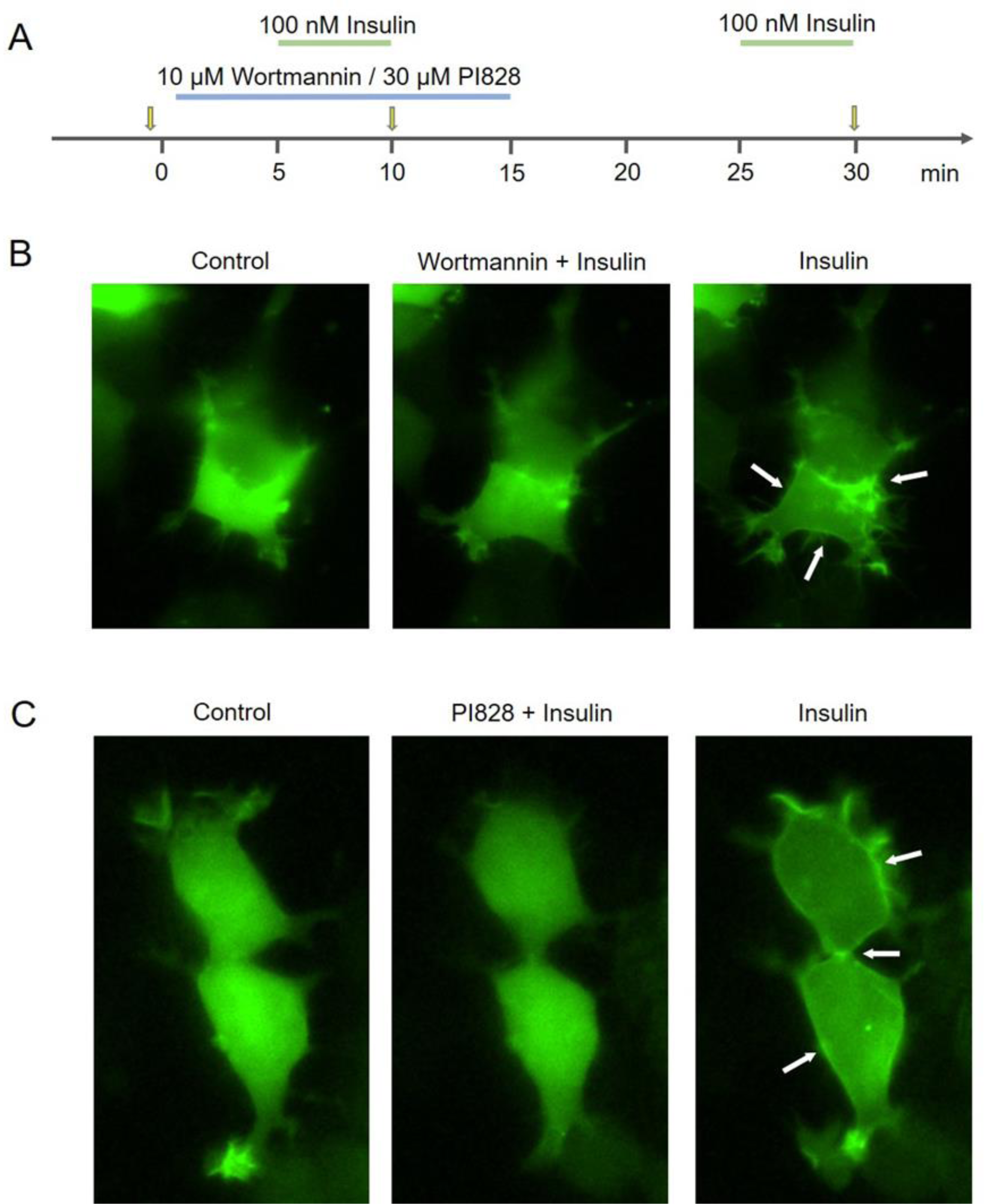
Evidence for the specific activity of wortmannin and PI828. (A) Experimental protocol timing. The applications of wortmannin, PI828, and insulin during recordings are indicated by the line segments above the time axis; the moments of image capturing are indicated by the arrows. (B, C) Representative sequential images of HEK/R-GECO1/PH(Akt)-Venus cells were obtained with SIM-microscopy at the moments indicated in (A). The left panels, control images of two cells obtained right before drug application demonstrate the virtually homogeneous distribution of PH(Akt)-Venus fluorescence over cell bodies. The middle panels, the cell stimulation with 100 nM insulin negligibly affected sensor distribution in the presence of 10 μM wortmannin (B) or 30 μM PI828 (C). The weak decrease in cell fluorescence was presumably arisen from photobleaching. The right panels, after the rinse of PI3K inhibitors, 100 nM insulin initiated a drop in the fluorescence of the cell cytosol and the appearance of detectable stripe-like fluorescent zones (arrows in the right panels), the phenomenon suggesting the insulin-induced accumulation of the PH(Akt)-Venus sensor in the plasmalemma.

### 3.3. Role of PI3K in ACh transduction

Since wortmannin was verified as an effective inhibitor of PI3K activity (Fig. 3B), its inability to affect cell responsiveness to aminergic agonists (Fig. 2) indicated that PI3K played a negligible role in their transduction. We verified this idea using HEK/R-GECO1/PH(Akt)-Venus cells, which provided the possibility of simultaneous monitoring intracellular Ca^2+^ and PI3K activity. These cells were stimulated by ACh and insulin in line with the protocol schematized in Fig. 4A. While in response to 1 μM ACh, cells massively generated Ca^2+^ transients lasting 60-100 s (Fig. 4B, upper panel), no evident translocation of the PH(Akt)-Venus sensor was observed (*n* = 846) (Fig. 4B, bottom panel), albeit ACh was present in the bath for nearly 10 min (Fig. 4A). Thus, ACh-induced Ca^2+^ signaling was not associated with the detectable stimulation of PI3K.

**Fig. 4.**
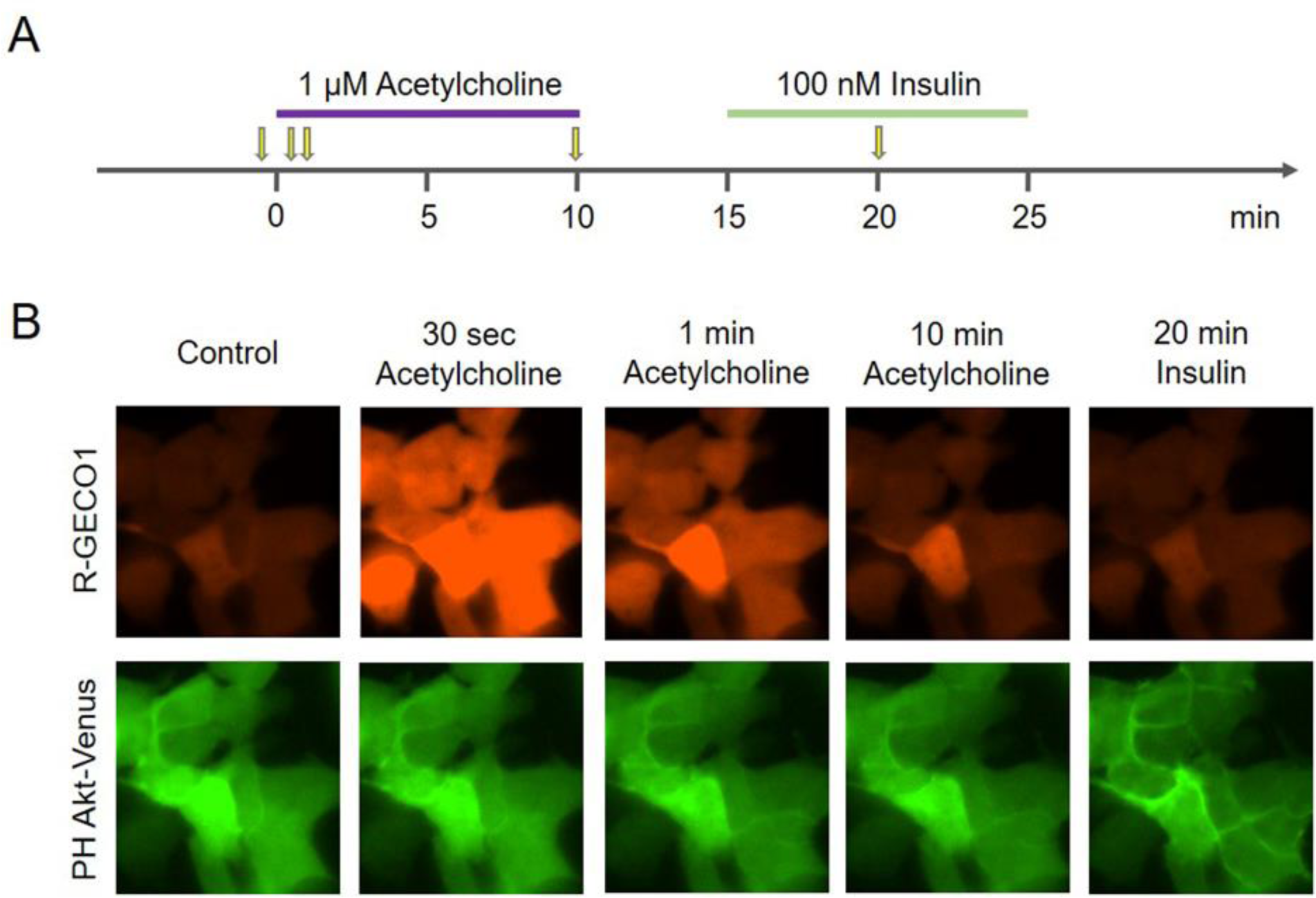
PI3K is not involved in ACh transduction. (A) Experimental protocol timing. The applications of ACh and insulin during recordings are indicated by the line segments above the time axis; the moments of image capturing are indicated by the arrows. (B) Representative sequential images of a group of HEK/R-GECO1/PH(Akt)-Venus cells obtained 30 s prior to ACh application (control) and after their stimulation by 1 μM ACh and 100 nM insulin. The upper and bottom panels represent the signals from the Ca^2+^ sensor and PIP_3_ sensor, respectively. As shown, 1 μM ACh elicited the transient increase in cytosolic Ca^2+^ but negligibly affected PI3K activity. In contrast, 100 nM insulin did not affect intracellular Ca^2+^ but stimulated PI3K activity.

At the same time, PI3K activity was rather inducible with 100 nM insulin in the same cells, judging by the translocation of PH(Akt)-Venus between their cytosol and plasmalemma, which occurred for 5 min after insulin application (Fig. 4B, bottom panel). It thus appeared that PI3K served neither as an effector downstream of muscarinic receptors nor as a modulator of ACh transduction machinery. In other words, the inhibition of PI3K could hardly underlie the collapse of ACh transduction in the presence of PI828 (Fig. 1A), and therefore this compound acted through a cellular target(s) unrelated to PI3K.

### 3.4. Evidence for an extracellular target of PI828

To elucidate a mechanism responsible for the effects of PI828 mentioned above, we focused on some details of Ca^2+^ mobilization triggered by the assayed agonists. It was particularly shown that the reduction of bath Ca^2+^ from 2 mM to 260 nM weakly or negligibly affected cellular responses to ACh (*n* = 163), histamine (*n* = 141), serotonin (*n* = 158), norepinephrine (*n* = 89), and ATP (*n* = 171), which however completely disappeared in the presence of inhibitor PLC U73122 (2 μM) (Fig. 5). These observations strongly argued that all used agonists mobilized Ca^2+^ largely by stimulating appropriate GPCRs coupled to the phosphoinositide cascade.

**Fig. 5.**
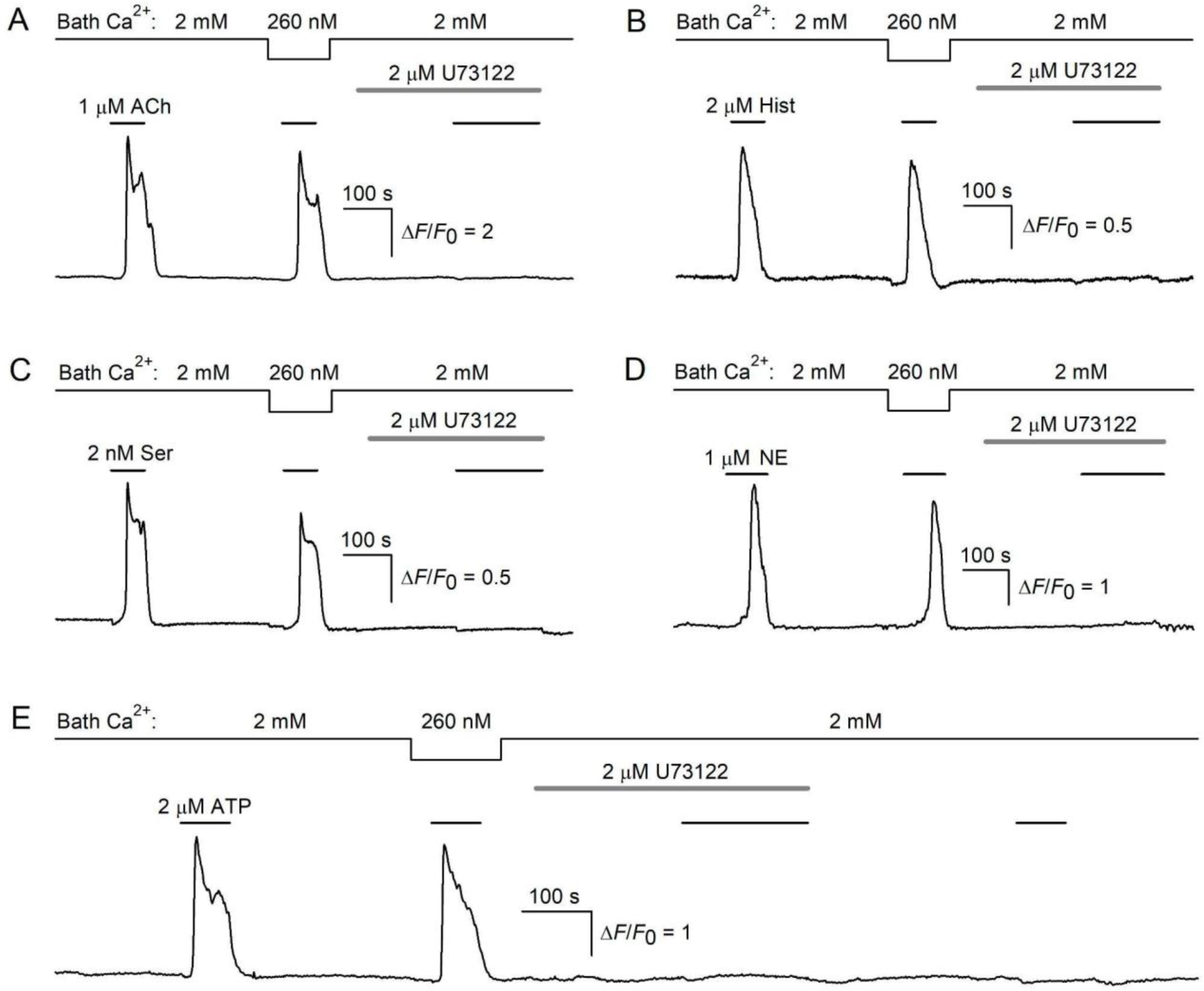
Involvement of the phosphoinositide cascade in aminergic and purinergic transduction. Ca^2+^ signals elicited by 1 μM ACh in HEK-293 cells (A), 2 μM histamine in C6 cells (B), 2 nM serotonin in CHO/5HT_2C_ cells (C), 1 μM norepinephrine (NE) in HEK-293 cells, 2 μM ATP in HEK-293 cells (E) were weakly affected by the reduction of bath Ca^2+^ from 2 mM to 260 nM and suppressed by the by PLC inhibitor U73122 (2 μM). The effects of U73122 were apparently irreversible (E).

Given that PI828 negligibly affected transduction of ATP and norepinephrine (Fig. 1), the phosphoinositide cascade as such could hardly be the main PI828 target. Otherwise, this compound would detectably influence Ca^2+^ responses to ATP and norepinephrine, as was the case with ACh, histamine, and serotonin (Fig. 1). The peculiar feature of the PI828 effects on ACh-, histamine-, and serotonin-responsive cells was that this PI3K inhibitor prevented agonist-induced Ca^2+^ mobilization even if it was applied simultaneously with the particular agonist (Fig. 1A–C, Fig. 2). On the other hand, the inhibition of intracellular enzymes by an extracellularly applied substance should have occurred with a certain delay because it would take some time for penetrating through the plasma membrane and being accumulated in the cytosol at an appropriate level. The fact that PI828 was capable of suppressing Ca^2+^ responses without preincubation pointed out the possibility that this chemical acted extracellularly. If so, the direct interaction of PI828 with GPCRs recognizing ACh, histamine, and serotonin might be the most likely way for the inhibition of agonist-induced signaling.

### 3.5. Muscarinic M3 receptor as a possible target of PI828

Among muscarinic receptors expressed in HEK-293 cells, the M3-receptor represents the dominant isoform (Atwood et al., 2011). Our consistent finding was that nanomolar doses of 4-DAMP, a specific antagonist of the muscarinic M3 receptor, invariably impaired Ca^2+^ responses of HEK-293 cells to ACh (n=162) (Fig. 6). Interestingly, the inhibitory effect of 4-DAMP developed quite rapidly, given that it was observed even if this antagonist was applied together with ACh (Fig. 6). The phenomenological similarity of the effects of PI828 (Fig. 1, Fig. 2) and 4-DAMP (Fig. 6) supported, albeit indirectly, the idea that PI828 could inhibit the responsiveness of cells to ACh by acting extracellularly on muscarinic receptors. In this case, the muscarinic M3 receptor appeared to be a likely extracellular target of PI828 in cholinergic HEK-293 cells.

**Fig. 6.**
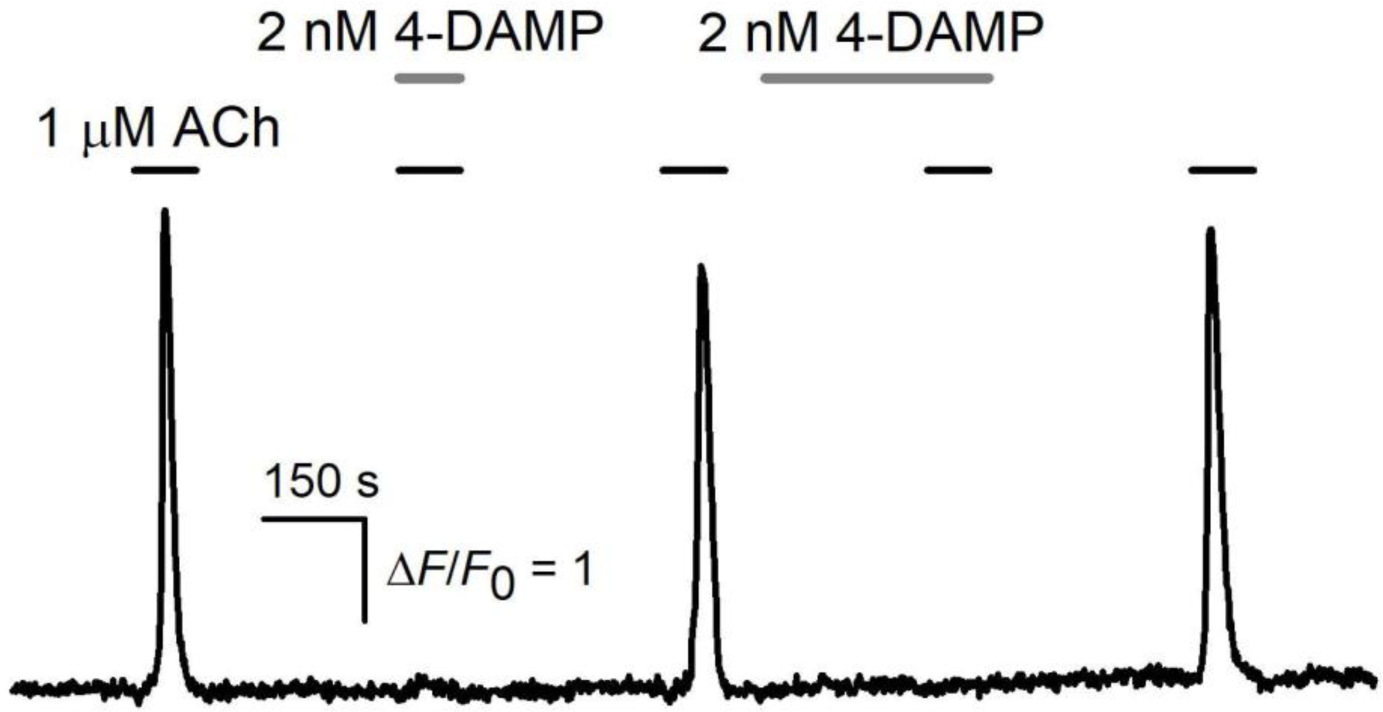
Representative monitoring of intracellular Ca^2+^ in a HEK-293 cell stimulated by ACh (1 μM) in control and in the presence of 4-DAMP (2 nM)

### 3.6. Computer modeling of interaction between PI828 and the muscarinic M3 receptor

As a way for verifying the direct interaction between PI828 and the muscarinic M3 receptor, we employed methods of computational biophysics. Molecular docking was involved in determining energetically most advantageous poses of the used compounds on the receptor surface. The stability of the ligand-receptor complexes predicted by docking was examined with molecular dynamics (MD).

Based on a number of available crystal structures of muscarinic receptors in complexes with the known antagonists (e.g. tiotropium, PDB ID: 4U15), the region suitable for docking was chosen to be bounded by transmembrane helices, extracellular loops, and the geometrical center of the muscarinic M3 receptor. In line with docking predictions, the most effective binding of ACh occurred deeply in the receptor transmembrane cavity (Fig. 7A). Since no 3D-structure of the ACh-M3 receptor complex (ACh-M3 complex) is currently available, the docking-determined location of ACh was considered as an appropriate fit for the orthosteric ACh-binding site at the muscarinic M3 receptor. The evolution of the docking-predicted ACh-M3 complex was then examined using the MD approach. A sample of three 100 ns MD trajectories was generated, and all of them showed that the ACh-M3 complex was quite stable for 100 ns at least (Fig. 7B). These findings indirectly supported the idea that the docking-predicted position of ACh at the M3-receptor fitted the orthosteric site properly (Fig. 7A).

**Fig. 7.**
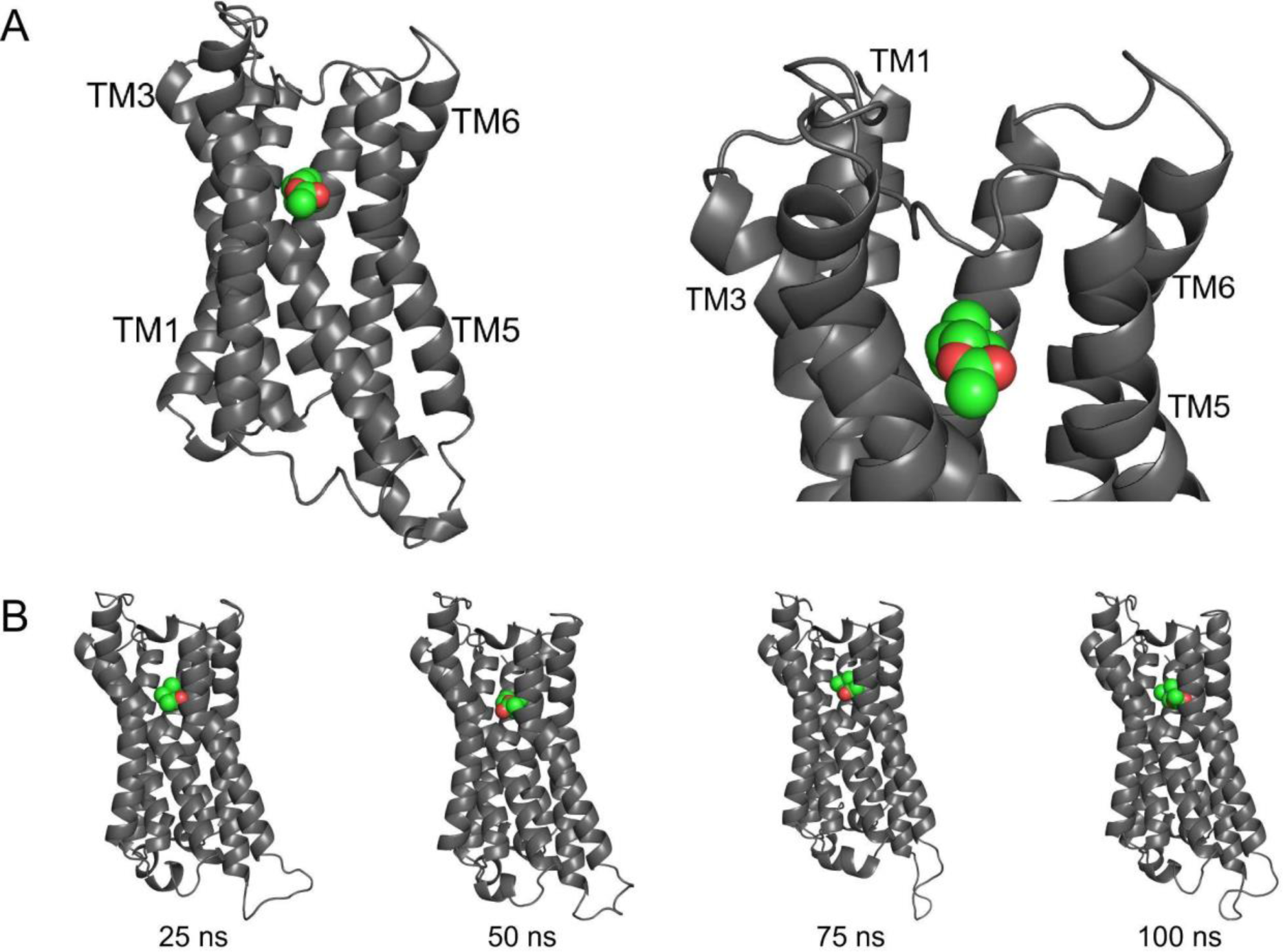
Computer modeling of muscarinic M3 receptor in complex with ACh. (A) Docking-predicted ACh-M3 complex, wherein ACh is shown as an assembly of colored spheres representing the carbon (green) and oxygen (red) atoms. TM1, TM3, TM5, and TM6 are 1st, 3rd, 5th, and 6th transmembrane helices, correspondingly. The left panel, the overall view of the ACh-M3 complex. The right panel, close-up view of the extracellular loops and ACh in the suggested orthosteric site. (B) Sequential snapshots taken from the representative MD trajectory at 25, 50, 75, and 100 ns. All MD trajectories, which started from the conformation shown in (A), indicated that ACh stably occupied the orthosteric site for 100 ns at least.

Next we analyzed interactions of the muscarinic M3 receptor with 4-DAMP, wortmannin, and PI828. Docking simulations showed that 4-DAMP was located in the orthosteric site (Fig. 8A). Consistently with the pharmacological activity as a M3 antagonist, 4-DAMP in this position should have prevented ACh binding. In contrast, wortmannin and PI828 were bound in the extracellular loop region, in the so-called vestibule (Fig. 8B,C). This raised the question why wortmannin and PI828 exhibited markedly different capabilities to inhibit ACh responses mediated by muscarinic M3 receptors (Fig. 2). Presumably, PI828 somehow interfered with the binding of ACh to the orthosteric site, while wortmannin was incapable of doing that, albeit both were bound in the vestibule (Fig. 8B,C). For instance, PI828 could occasionally enter into the orthosteric site from the vestibule, while wortmannin could solely dissociate to the extracellular space. We therefore performed MD simulations to determine evolution of the docking-predicted complexes of the muscarinic M3 receptor with 4-DAMP, wortmannin, and PI828.

**Fig. 8.**
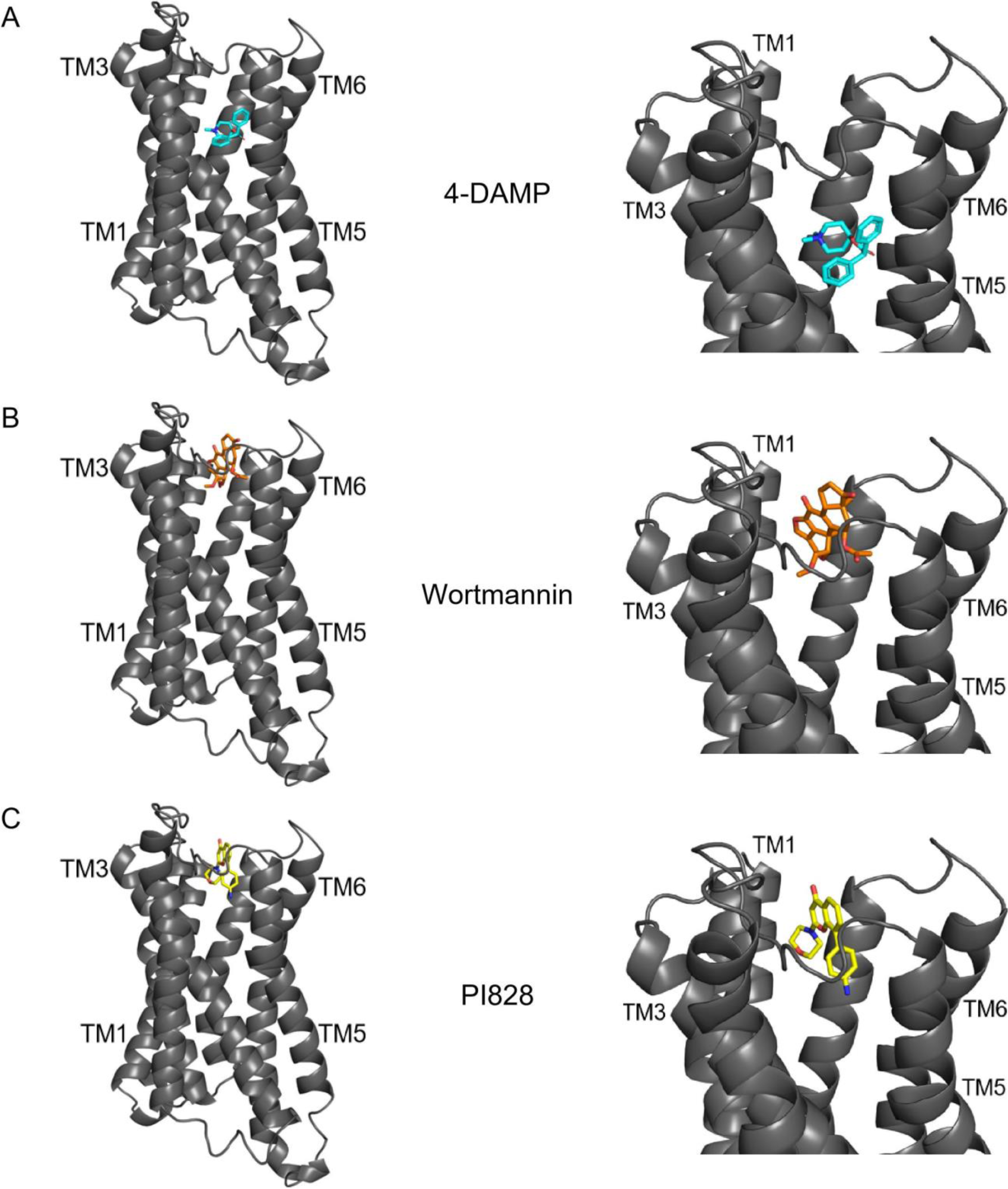
Docking-predicted conformations of the muscarinic M3 receptor in complex with 4-DAMP (A), wortmannin (B), PI828 (C). The left panels represent the overall view of the receptor; the right panels illustrate the extracellular loop region and the orthosteric site. TM1, TM3, TM5, and TM6 are as in Fig.7A. As shown, 4-DAMP localizes in the orthosteric site, while wortmannin and PI828 occupy the extracellular loops region.

From a sample of generated trajectories it turned out 4-DAMP was retained in its position in the orthosteric site (Fig. 9A) for 100 ns at least, thus preventing ACh binding and M3-receptor activation. As shown previously, antagonists of the muscarinic receptors, particularly tiotropium, could briefly occupy the receptor vestibule prior to entering into the orthosteric site of the receptor (Kruse et al., 2012). Similar simulations showed that being initially bound in the vestibule (Fig. 9B), wortmannin tended neither to come into the orthosteric site nor to leave the vestibule during 100 ns simulations. Given however that these standard MD trajectories were not sufficiently long, we examined the evolution of the wortmannin-M3 receptor complex by using the accelerated MD approach (see Amber manual). Even during these effectively much more prolonged simulations, wortmannin did not proceed to the orthosteric site or dissociate to the extracellular space. Interestingly, some of these accelerated simulations raised the possibility that even with wortmannin bound, there could be enough space in the vestibule to allow ACh to pass through the vestibule toward the orthosteric site (Fig. 10). This would reconcile the simulation results, which suggested specific binding of wortmannin to the muscarinic M3 receptor (Fig. 9), and experimental findings validating its inability to inhibit ACh responses (Fig. 2). Much longer simulations are required to clarify this issue.

**Fig. 9.**
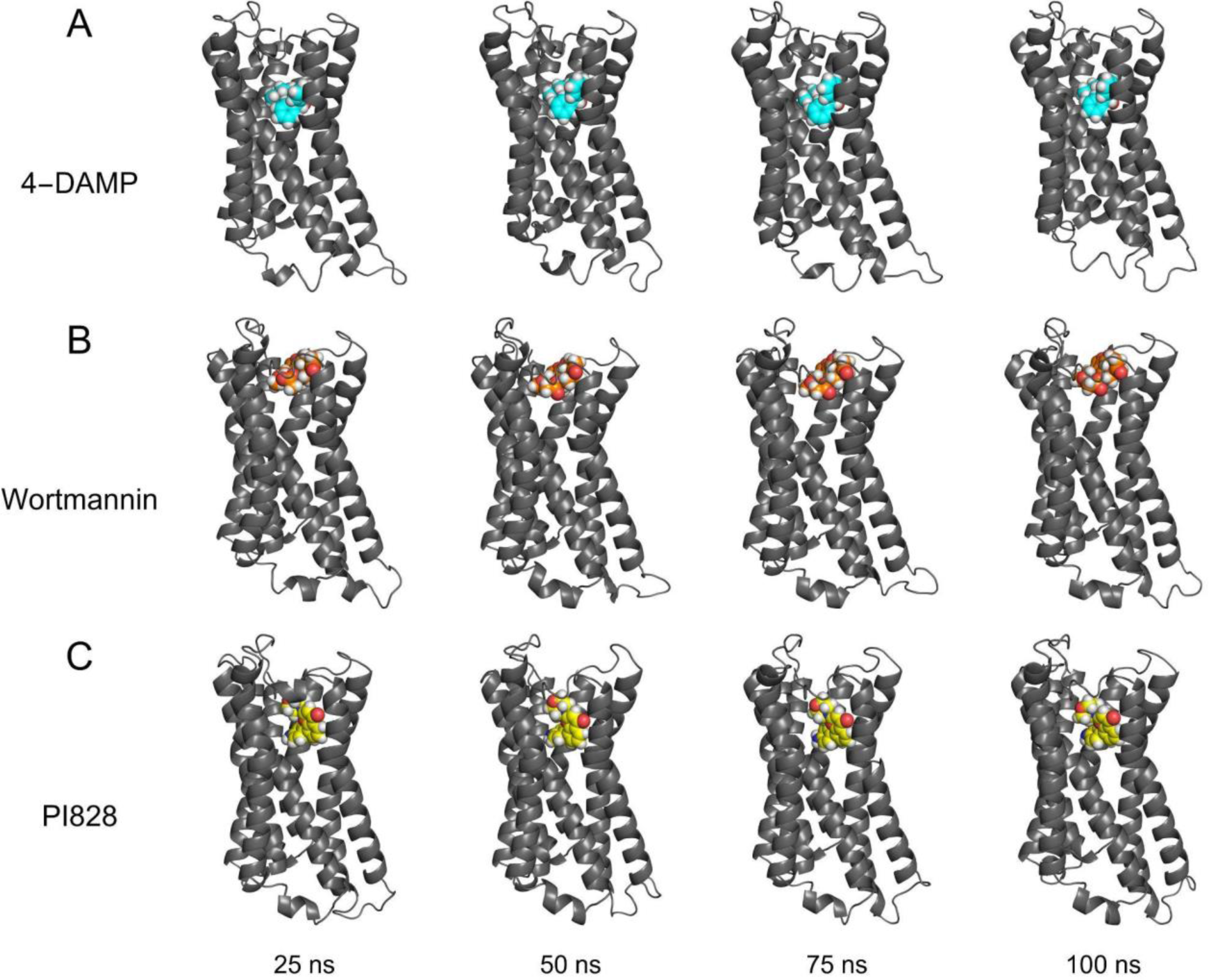
Muscarinic M3 receptor in complex with the studied compounds modeled with MD simulations. Snapshots are taken from representative MD trajectories at 25, 50, 75, and 100 ns. The MD simulations for 4-DAMP (A), wortmannin (B), and PI828 (C), started from the conformations shown in Fig. 8A,B,C, respectively. The compounds are shown as spheres colored according to element (carbon – cyan, orange, and yellow in 4-DAMP, wortmannin, and PI828, correspondingly; nitrogen – blue; oxygen – red; hydrogen – white). As demonstrated, 4-DAMP and wortmannin did not leave the orthosteric site and the vestibule, respectively, within 100 ns. At the same time, PI828 was capable of moving from its docking-predicted location in the vestibule towards the orthosteric site already after 25 ns.

**Fig. 10.**
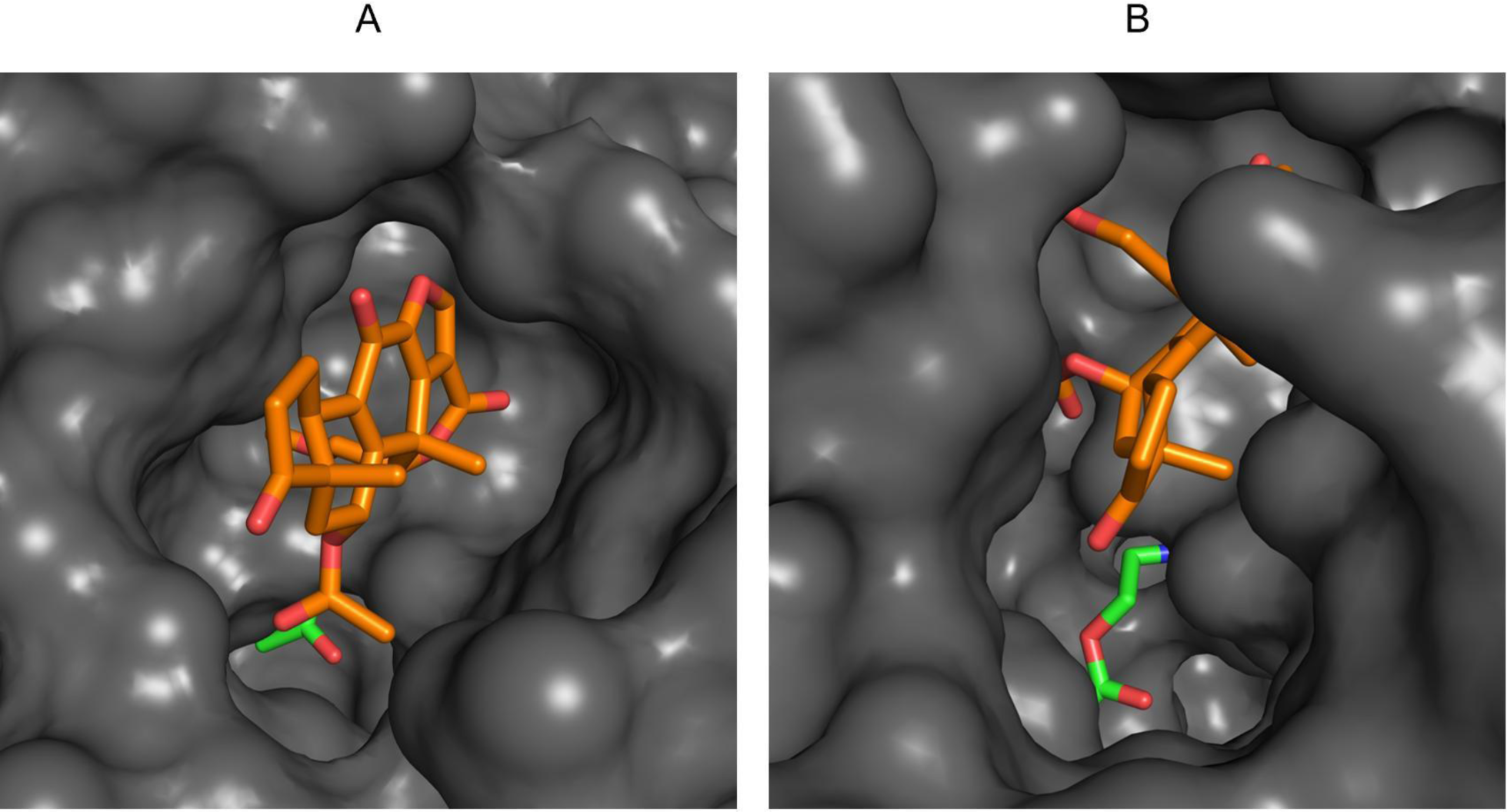
Wortmannin positions in the vestibule of the muscarinic M3 receptor seen from the extracellular solution. Conformations of the muscarinic M3 receptor in complex with wortmannin (orange), here ACh (green) in its position at the orthosteric site of the receptor is shown for reference. (A) Docking-predicted conformation of the complex. In this conformation wortmannin would apparently prevent ACh crossing from the extracellular solution through the vestibule to occupy the orthosteric site. (B) Conformation obtained from the accelerated MD simulation. Wortmannin can be seen to be displaced, presumably leaving enough space in the vestibule for ACh to pass.

The 100 ns MD simulations of the muscarinic M3 receptor in complex with PI828 revealed that PI828 could indeed leave the initial docking-predicted position in the vestibule and move toward the orthosteric site (Fig. 9C, Fig. 11 A,B). For the particular trajectory shown in Fig. 11, the initial distance between geometric centers of ACh and PI828, taken as average coordinates of their heavy atoms, of 11.27 Å (Fig. 11A) decreased to 4.43 Å (Fig. 11B). As suggested previously, the tyrosine lid, which comprises tyrosines 148, 506, and 529, separates the vestibule from the orthosteric site (Kruse et al., 2012). In our simulations, terminal oxygen atoms displaced by 4.30 Å in Tyr 148 and by 1.66 Å in Tyr 506 (Fig. 11C), and these conformational transformations presumably opened a passage for PI828 toward the orthosteric site.

**Fig. 11.**
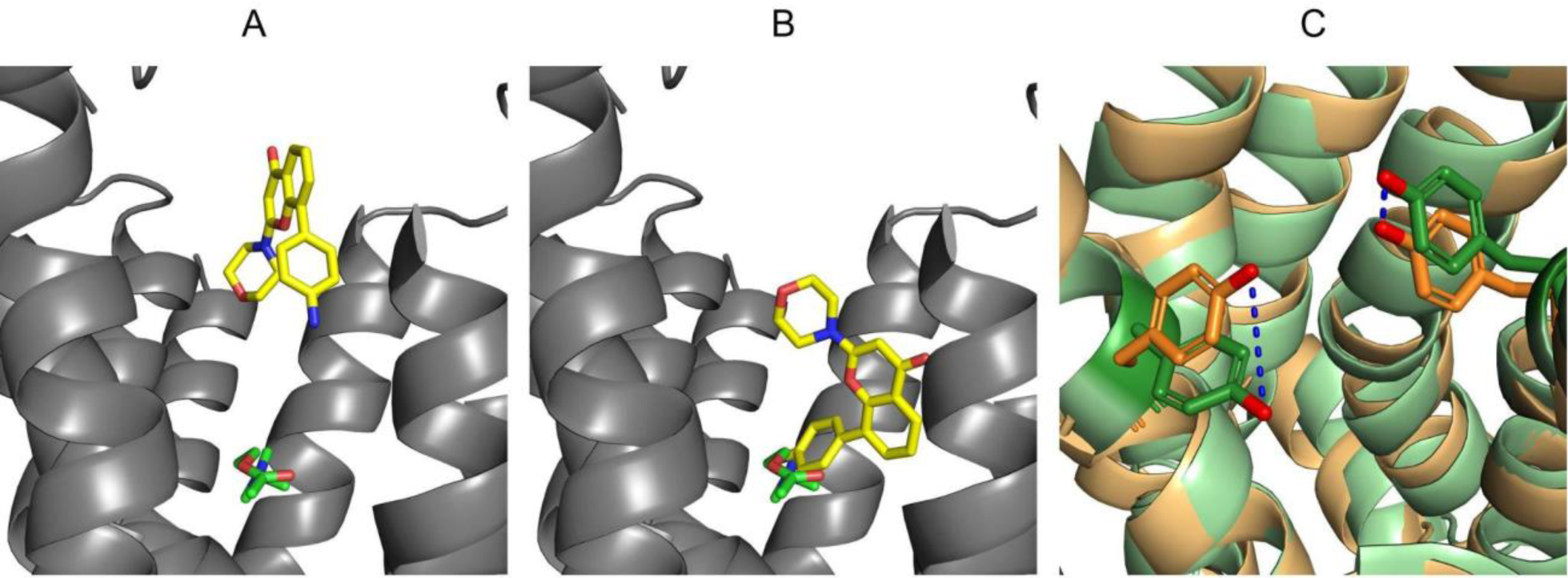
Passage of PI828 from the M3 receptor vestibule into its orthosteric site. Snapshots of PI828-M3 receptor complex obtained with representative MD-simulation. Positions of PI828 (yellow) in the receptor vestibule predicted by docking (A) and obtained from over 100 ns-long trajectory showing its passage toward the orthosteric site (B); here ACh (green) is shown in its position at the orthosteric site of the receptor for reference. (C) Transformation of the tyrosine lid opening the passage for PI828. Tyrosines 148 and 506 are rendered as sticks, the muscarinic M3 receptor in docking-predicted conformation is shown in orange and after molecular dynamics simulation – in green. Note the displacement of the tyrosine terminal oxygen atoms shown in red.

Finally, computer modeling showed that 4-DAMP and PI828 actually could compete with ACh for the orthosteric site, thus preventing activation of the muscarinic M3 receptor. By occupying the vestibule, wortmannin could potentially block it and preclude ACh entry to the orthosteric site. Nevertheless, our simulations suggested the existence of such a conformation of the wortmannin-M3 receptor complex, which could allow ACh to pass the vestibule and activate the receptor. These computational findings were consistent with the experimental results and plausibly explained why PI828 was capable of suppressing responses of HEK-293 cells to ACh, while wortmannin was ineffective as a M3 receptor antagonist.

## 4. Discussion

The first identified PI3K inhibitors were wortmannin (Arcaro and Wymann, 1993; Yano et al., 1993), a naturally occurring metabolite of *Penicillium funiculosum*, and LY294002 (Vlahos et al., 1994), which was derived from the flavonoid quercetin. Being considered as rather specific PI3K inhibitors for a long time, wortmannin and LY294002 were widely used to verify a role for the PI3K pathway in cellular functions. However, these PI3K inhibitors were later shown not only to impair activity of PI3Ks from different classes but also to affect unrelated proteins involved in various physiological processes, particularly, metabolism, transcription or protein trafficking and dynamics (Gharbi et al., 2007). Notable among them are non-lipid kinases, such as mTOR (Brunn et al., 1996), DNA-dependent protein kinase (Smith et al., 1999), CK2 (Davies et al., 2000), Pim-1 kinase (Jacobs et al., 2005), nuclear factor-κB (Choi et al., 2004; Kim et al., 2005), MRP1 (Abdul-Ghani et al., 2006). In studies of intracellular signaling, wherein both wortmannin and LY294002 were used in parallel, dramatic differences in their effects has been documented. For instance, it was shown that LY294002, but not wortmannin, irreversibly inhibited Ca^2+^ responses to carbachol and histamine in bovine and human airway smooth muscle (ASM) cells (Ethier and Madison, 2002). In this study, LY294002 (50 μM) alone increased cytosolic Ca^2+^ concentration by mobilizing intracellular calcium stores, and this mobilization was not canceled by the PLC inhibitor U73122 and IP_3_ receptor antagonist xestospongin C. Although a related target was not identified with confidence, the authors inferred that LY294002 depleted Ca^2+^ store, thereby rendering cells incapable of generating Ca^2+^ signals to carbachol and histamine. In rat ASM cells, wortmannin and LY294002 had opposite effects on serotonin-induced Ca^2+^ signals: while wortmannin caused a small increase in Ca^2+^ responses, LY294002 significantly decreased their magnitudes (Ethier and Madison, 2002). Based predominantly on biochemical evidence, the authors inferred that LY294002 affected the serotonin responses independently of PI3K suppression, by acting on, or upstream of, PLC, possibly by inhibiting CK2.

The more potent PI3K antagonist PI828, an analog of LY294002 developed in search for its intracellular targets, also has been reported to cause nonspecific effects (Gharbi et al., 2007). In our study, PI828 reversibly suppressed Ca^2+^ transients elicited by some aminergic agonists, while wortmannin did not do so (Fig. 2). Given that at the used concentrations of 10 – 50 μM PI828 never triggered a thapsigargin-like increase in intracellular Ca^2+^ (Fig. 1), it was unlikely that the PI828 effects were associated with Ca^2+^ store depletion. Yet, PLC inhibition could not underlie the inhibitory effects of PI828 on cell response to ACh, serotonin, and histamine (Fig. 1), given that this compound negligibly affected Ca^2+^ transients elicited by ATP and norepinephrine through stimulation of the phosphoinositide cascade (Fig. 5). We also excluded the possibility that PI828 suppressed Ca^2+^ responses to ACh by acting on CK2, since unlike PI828, the CK2 inhibitor hematein undetectably affected ACh responsivity of HEK-293 cells (*n* = 56) (Fig. 12). Thus, at least in the given case, the inhibitory effects of PI828 were not associated with the CK2 inhibition.

**Fig. 12.**
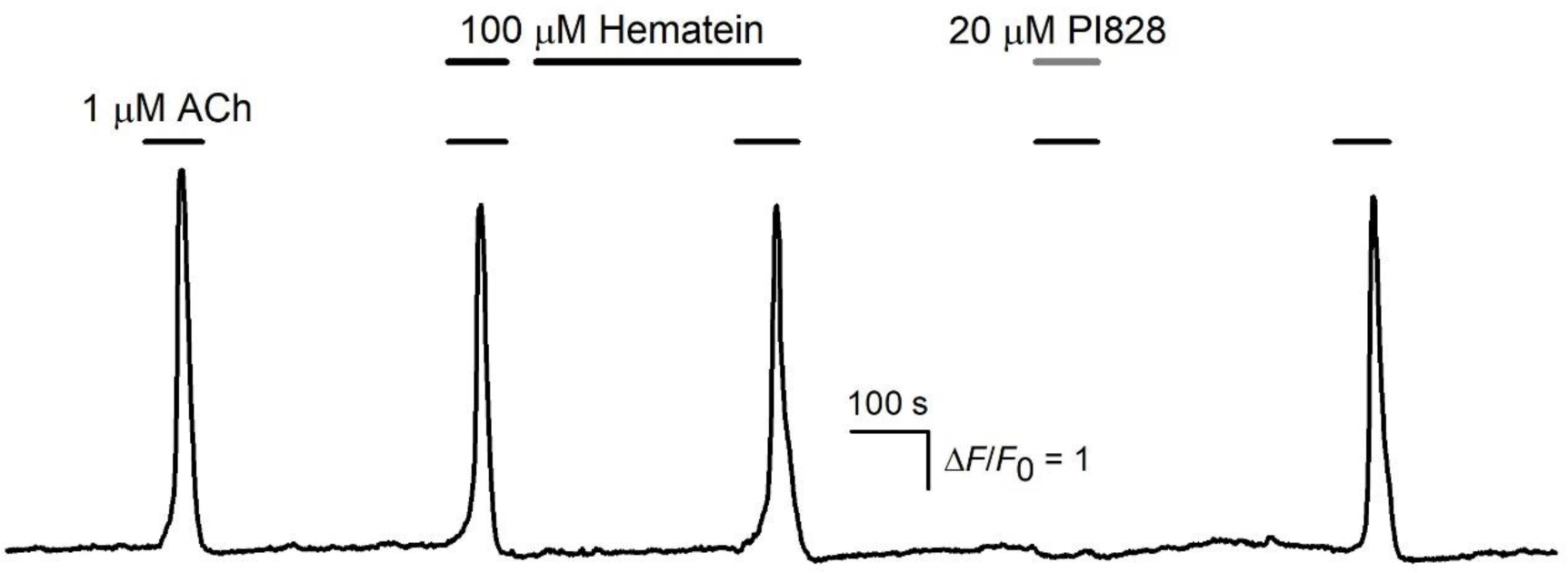
Effects of CK2 inhibitor and PI828 on cells responsiveness to ACh. Representative Ca^2+^ transients elicited by 1 μM ACh in HEK-293 cells in control and in the presence of hemantein (100 μM) or PI828 (20 μM).

In the previous study (Kotova et al., 2020), we showed that LY294002 and its structural analogue LY303511 impaired Ca^2+^ signaling induced by histamine, ACh, and serotonin in human mesenchymal stromal cells and in CHO/5-HT_2C_ cells. The obtained evidence raised the possibility that LY294002 and LY303511 could directly affect activity of aminergic GPCRs involved. Interestingly, based on the similarity of the structural nuclei of LY294002 and ketanserin, a classic 5-HT_2A_ receptor antagonist, it was suggested in 1999 that LY294002 might serve as an inhibitor of serotonin receptors (Banes et al., 1999). Moreover, the inhibitory activity of ketanserin against H1-histamine and alpha1-adrenergic receptors has also been documented (Marin et al., 1990). We adopted the idea about the correlation between the structural similarity of LY294002 and ketanserin and their functional activities, and surmised that it also could be applicable to the PI828/ketanserin pair (Fig. 13A). Amazingly, ketanserin was indeed capable of inhibiting Ca^2+^ responses of HEK-293 to ACh in a PI828-like manner (*n* = 82) (Fig. 13 B). Specifically, ketanserin and PI828 inhibited Ca^2+^ responses to ACh at the similar IC_50_ of about 20 μM, and both were effective being applied concurrently with the agonist.

**Fig. 13.**
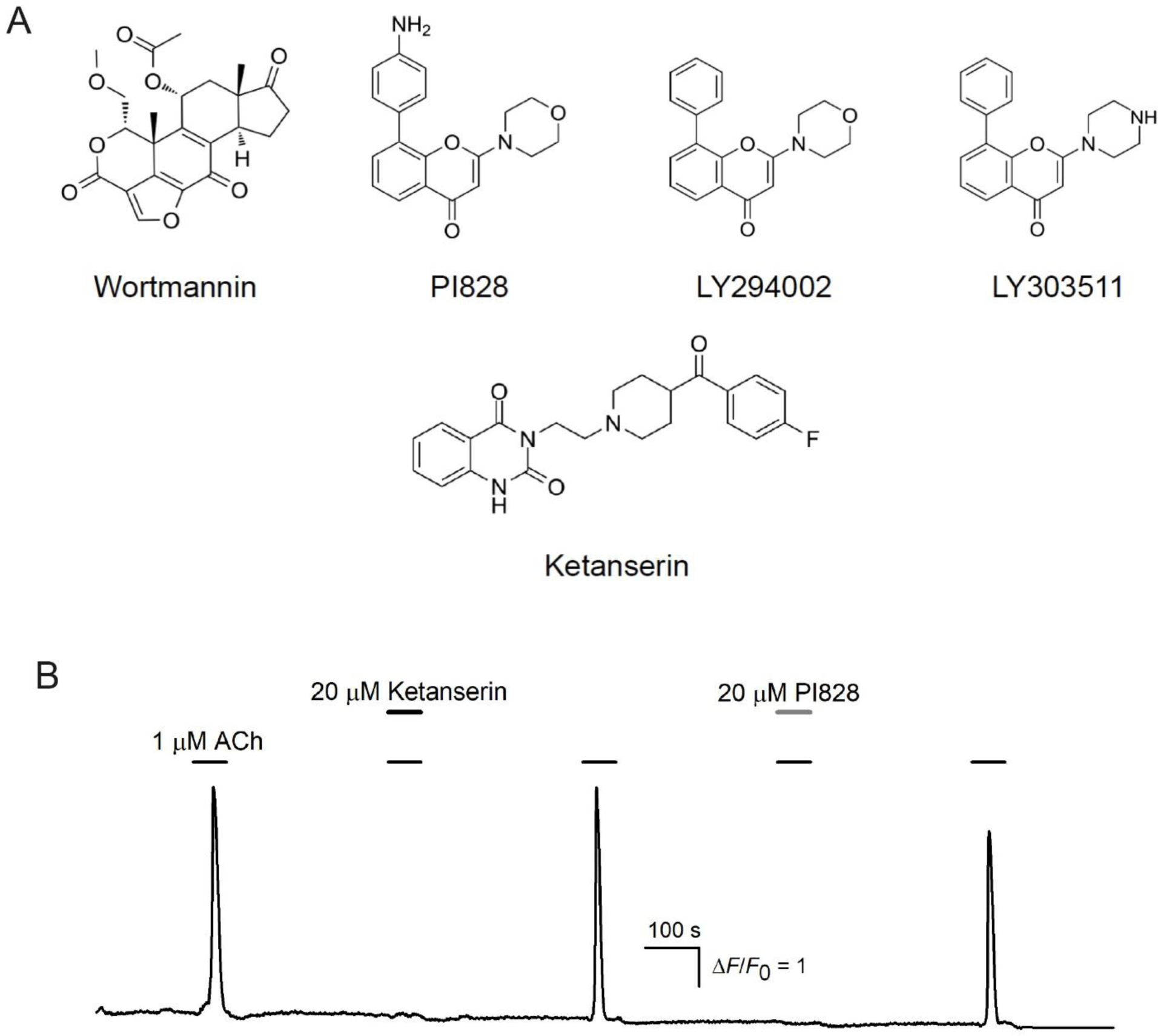
PI3K inhibitors and ketanserin. (A) Structural formulas of PI3K inhibitors and ketanserin. (B) Representative Ca^2+^ transients elicited by 1 μM ACh in HEK-293 cells in control and in the presence of ketanserin (20 μM) or PI828 (20 μM).

To summarize, the previous data obtained by us and others clearly indicated that intracellular Ca^2+^ mobilization initiated by aminergic agonists could be suppressed by LY294002 and LY303511 by means of mechanisms unrelated to PI3K inhibition (Banes et al., 1999; Ethier and Madison, 2002; Kotova et al., 2020; Tolloczko et al., 2004). Consistently, stimulation of cells with serotonin and histamine did not lead to activation of Akt and therefore PI3K (Alonso et al., 2016; Tolloczko et al., 2004). In the present study, we showed that PI828, one more compound developed initially as a PI3K inhibitor, completely and reversibly suppressed Ca^2+^ signaling initiated by ACh, histamine, and serotonin (Fig. 1, 2). Using genetically encoded sensors for PIP_3_, we demonstrated that ACh did not stimulate PI3K activation and PIP_3_ generation in HEK-293 cells responsive to the agonist with Ca^2+^ mobilization (Fig. 4). Taken together, the data of our physiological experiments, reported previously and presented here, led us to the conclusion that LY294002, LY303511, and PI828 suppressed cellular responses to certain aminergic agonist most likely by acting as antagonists of the corresponding GPCRs. Computer simulations supported the possibility that this mechanism could be feasible (Figs. 7–11) (Kotova et al., 2020).

## 5. Conclusions

Here we confirmed at the level of individual cells that wortmannin and PI828 are effective PI3K inhibitors and that GPCR signaling does not necessarily recruit PI3K. We also showed that PI828 suppressed Ca^2+^ signaling initiated by ACh, histamine, and serotonin, and that the underlying mechanism was not related to the inhibition of PI3K activity. Most likely, PI828 acted directly on aminergic GPCRs involved as an antagonist. Thus, PI828 should be used cautiously to demonstrate PI3K contribution to GPCR signaling.

## CRediT authorship contribution statement

**Polina D. Kotova:** Supervision, Investigation, Visualization, Writing – original draft, Funding acquisition. **Ekaterina A. Dymova:** Investigation, Visualization. **Oleg O. Lyamin:** Investigation, Formal analysis, Visualization, Writing – original draft. **Olga A. Rogachevskaja:** Investigation. **Stanislav S. Kolesnikov:** Conceptualization, Writing – review & editing

## Acknowledgements

The authors thank Daria M. Potashnikova for her assistance in sorting cells on the FACSAria SORP cell sorter (MSU Program of Development).

